# No Electrophysiological Evidence for Semantic Processing During Inattentional Blindness

**DOI:** 10.1101/2024.05.31.596114

**Authors:** Brendan T. Hutchinson, Bradley N. Jack, Kristen Pammer, Enriqueta Canseco-Gonzalez, Michael Pitts

**Affiliations:** Faculty of Behavioural and Movement Sciences, Vrij University; Research School of Psychology, Australian National University; School of Psychology, University of Newcastle; School of Psychology, Reed College

**Keywords:** perception, N400, inattentional blindness, attention, semantic, consciousness

## Abstract

A long-standing question concerns whether sensory input can reach semantic stages of processing in the absence of attention and awareness. Here, we examine whether the N400, an event related potential associated with semantic processing, can occur under conditions of inattentional blindness. By employing a novel three-phase inattentional blindness paradigm designed to maximise the opportunity for detecting an N400, we found no evidence for it when participants were inattentionally blind to the eliciting stimuli (related and unrelated word pairs). In contrast, when minimal attention was allocated to the same task-irrelevant word pairs, participants became aware of them, and a small N400 became evident. Finally, when the same stimuli were fully attended and relevant to the task, a robust N400 was observed. In addition to univariate ERP measures, multivariate decoding analyses were unable to classify related versus unrelated word pairs when observers were inattentionally blind to the words, with decoding reaching above-chance levels only when the words were (at least minimally) attended. Our results also replicated several previous studies by finding a “visual awareness negativity” (VAN) that distinguished task-irrelevant stimuli that were perceived compared with those that were not perceived, and a P3b (or “late positivity”) that was evident only when the stimuli were perceived and task relevant. Together, our findings suggest that semantic processing might require at least a minimal amount of attention.

## Introduction

The visual world is processed to a remarkable extent in the absence of awareness. Low-level features such as colour and basic shape are processed irrespective of an observer’s ability to report that they perceived those features (Ro, Singhal, Breitmeyer, & Garcia, 2009). Unconscious processing continues up to intermediate stages of the visual hierarchy, such as surface-boundary integration (Guzzon & Casco, 2011) and the grouping and segregation of features (Pitts, Martínez, & Hillyard, 2012; Scholte, Witteveen, Spekreijse, & Lamme, 2006; Schelonka, Graulty, Canseco-Gonzalez, & Pitts, 2017). Even perceptual inference-like processes, such as depth perception and illusory contour formation, seemingly occur in the absence of awareness (Fahrenfort et al., 2017; Vandenbroucke et al., 2014). With the brain’s capacity to extract and process so much visual information unconsciously, one may wonder what is left that requires conscious awareness.

An unresolved question is whether semantic information can be processed in the absence of awareness (Deacon & Shelley-Tremblay, 2000; Mudrik & Deouell, 2022). While much research has suggested fairly robust unconscious semantic processing (Deacon et al., 2000; Grossi, 2006; Kiefer, 2002; Luck et al., 1996; van Gaal et a., 2014), previously reported positive findings are highly contingent on the methods used to manipulate awareness (Batterink, Karns, Yamada, & Neville, 2009; Damian, 2001; Davis et al., 2007; Holcomb & Grainger, 2009; Holcomb, Reder, Misra, & Grainger, 2005). For example, with some awareness manipulations, visual attention can still be allocated to the (unseen) stimuli (Dehaene et al., 1998; Naccache et al., 2002; Schnuerch et al., 2016), while with other manipulations, attention to the stimuli is disrupted (Batterink et al., 2010). In the current study, we contribute novel evidence to this area by examining whether the N400—a classic event related potential (ERP) component associated with semantic processing—can occur under conditions of inattentional blindness.

### The N400

The N400 is one of the most reliable ERPs studied (Kutas & Federmeier, 2011). Named for its negative-going peak that occurs at approximately 400 milliseconds post stimulus, the N400 is characterised as a robust slow-wave, commonly observed over bilateral central-posterior regions, and is elicited when a previously established semantic context is violated. The N400 was first reported in the pioneering work of Kutas and Hillyard (1980) in a variant of the classic oddball paradigm, where the congruency of the closing word of a sentence was varied with respect to the context established by the sentence. For example, an N400 was evident when the closing word of the sentence “the woman drove to work in her shiny new *nose*” was compared with “the woman drove to work in her shiny new *car*” (Kutas & Hillyard, 1980). It was quickly discovered that this effect could be obtained with other stimuli, such as pictures and sounds, so long as they established a semantic context that could subsequently be violated. Since then, it has been found that the N400 is also elicited by simple pairs of words (Holcomb & Grainger, 2009). For example, an N400 is elicited by the second of two sequentially presented words (referred to as the *target*) if it is incongruent with the semantic context established by the first word (referred to as the *prime*), such as Doctor-Nurse compared with Apple-Nurse (McCarthy & Nobre, 1993).

The N400 is considered a reliable index of semantic processing and is commonly used as an outcome measure for exploring the degree to which semantic processing can occur in different task situations, such as when attention is distracted and stimuli are processed unconsciously or “automatically” (Holcomb et al., 2005; for review, see Kutas & Federmeier, 2011). To date, results have been inconclusive. Some studies have found the N400 is elicited even when observers are not aware of the eliciting stimuli (Kiefer, 2002). In masking studies, for example, the N400 persists when the target is masked (Stenberg, Lindgren, Johansson, Olsson, & Rosén, 2000) and sometimes if the prime is masked (Deacon et al., 2000; Grossi, 2006, but see Holcomb & Grainger, 2009; Van Gaal et al 2014). More recent masking studies suggest that unconscious N400 effects may depend on aspects of the task design, such as the distance between the prime and target. Using word primes and picture targets, Mongelli et al. (2019) varied the temporal position between prime and target and found an N400 for masked words as primes, but only when they were presented at the closest temporal position to the target picture. A similar finding was made by Nakamura et al. (2018), who reported evidence for N400 only if the distance between words did not exceed two words.

Work using other manipulations of awareness suggest that the N400 is observed if the target is unattended, but not if the prime is unattended. Early influential work on the attentional blink found that the N400 was elicited for “blinked” target stimuli—that is, for stimuli participants were unable to report as perceived (Luck et al., 1996). However, these findings have been challenged by research that found no N400 for masked primes (Holcomb et al., 2005; Holcomb & Grainger, 2009) or for incorrectly reported target stimuli during the attentional blink (Batterink et al., 2010). Also, the N400 is abolished when stimuli are rendered unconscious due to continuous flash suppression or binocular rivalry (Kang, Blake, & Woodman, 2011; also see Sklar et al., 2012). From a broad perspective, then, whether an N400 can be elicited in the absence of awareness remains unclear.

### Awareness, Attention and Task Relevance

One key design feature in many of these previous studies that has often been overlooked is that the stimuli employed were typically task relevant. Observers were—in almost all instances—explicitly required to respond to the N400 eliciting stimuli (Luck et al., 1996; Naccache, Blandin, & Dehaene, 2002). While studies have examined the N400 for task irrelevant stimuli (Erlbeck, Kubler, Kotchoubey, & Veser 2014; Neumann & Schweinberger, 2008; Relander, Rämä, & Kujala, 2008), task relevance and awareness have typically co-varied in such designs. Early work by McCarthy and Nobre (1993) found that the N400 was absent for stimuli that were task irrelevant and required no response. Yet this study did not employ a measure of whether participants consciously perceived the task irrelevant stimuli. Hence, it is not clear whether the effect was a result of (some combination of) whether the stimuli were perceived or task irrelevant.

The broader issue is that attention, task relevancy, and awareness may have been confounded in previous work examining the relation between semantic processing and awareness. This is particularly important because each of these factors may have unique effects on the N400. For example, Erlbeck and colleagues (2014) found that task instructions strongly attenuated the N400. In their study, participants were instructed to either explicitly attend to, passively ignore, or actively ignore auditory sentences. In the explicit attention condition, N400-eliciting stimuli were task relevant, whereas in the active ignore condition, participants attended to an alternative task and hence the eliciting stimuli were task irrelevant, with the passive ignore condition falling somewhere in between. The N400 was robust in the explicit attention condition, markedly reduced in the passive ignore condition, and entirely absent in the active ignore condition (Erlbeck et al., 2014). In more recent work, Szalardy et al. (2018) manipulated both attention and task relevance of auditory stimuli (speech streams) and found that both task relevant and task irrelevant syntactic violations elicited an N400 within the attended stream, but not the unattended stimulus stream. In other words, an N400 was elicited irrespective of task relevancy, but only when attended.

Conflating task relevancy and awareness is problematic for several reasons, most notably because ERP markers of visual consciousness vary with task relevancy (Cohen et al., 2020; Dellert et al., 2022). The most striking illustration of this concerns the P3b (or “late positivity”). Despite being considered one of the most reliable ERP components associated with consciousness, recent work suggests the P3b to be absent in response to clearly perceived but task irrelevant visual information. When awareness is manipulated independent of the task, an earlier and more subtle ERP (the “visual awareness negativity”, or VAN) has been found to index awareness of information (Pitts et al., 2012). Under such conditions, the P3b is not generated until the information becomes task relevant (Pitts et al., 2012; Pitts et al., 2014).

### Inattentional Blindness

Inattentional blindness (IB) refers to the failure to notice an unexpected stimulus when attention is engaged in a separate demanding task, despite the stimulus being within one’s visual field and otherwise clearly perceptible (for review, see Hutchinson, 2019, and Hutchinson et al., 2022). Surprisingly, no prior work has sought to assess whether the N400 can be elicited under conditions of IB. This is likely due to limitations with studying IB using electroencephalography (EEG). Experimental work on IB often includes only a single critical trial, because once the observer is questioned and thus cued-in to the existence of the critical stimulus, IB is far less likely to occur. Since multiple trials are needed to achieve adequate signal-to-noise ratios in neural data, this creates a problem for gaining sufficient data for electrophysiological analysis during IB. Recent work has overcome this issue through either modification of classic paradigms (e.g., Hutchinson et al., 2023) or through block-wise classification of data via retrospective questioning (e.g., Pitts et al., 2012). For example, in Pitts et al. (2012) three-phase paradigm, participants progressed through many trials of the task (where hundreds of critical stimuli were presented) before they were eventually questioned on whether they perceived the unexpected critical stimuli in phase one (the first block of trials). About half of the participants were deemed inattentionally blind. After being questioned (and thus cued to expect these stimuli), participants then completed another round of the same task on the same stimuli in phase two (here, all participants were aware of the still-task-irrelevant stimuli). Finally, in phase three, while the stimuli remained the same, the task was changed to render the critical stimuli task relevant. It was this methodology that led to the finding that the P3b is absent for clearly perceived but task irrelevant stimuli, while the earlier VAN distinguishes between perceived and not-perceived stimuli, even if they remain task irrelevant (Förster et al., 2020; Pitts et al., 2012; Dembski et al., 2021).

Whether semantic processing can occur during IB has been previously explored using behavioral measures. For example, both Schnuerch, Kreitz, Gibbons, and Memmert (2016) and Pugnaghi, Memmert, and Kreitz (2020) reported behavioural evidence for semantic processing of numbers during inattentional blindness. In their work, participants were to discriminate whether target numerals were smaller or larger than five, while unexpected number stimuli were repeatedly presented within the attended region of space. If semantic processing occurred in the absence of awareness, then performance on the target categorisation task should have been hindered when the unexpected numbers conflicted with the category of the target numeral (for example, if the target number was six and the unexpected stimulus number was four). In fact, this is what was found, as reaction times were slightly (but consistently) slower when the unexpected stimuli were incongruent with the target item (but see Pugnaghi et al., 2020, experiment 2).

Other recent work employed EEG to examine the extent to which more basic types of linguistic processing can occur during IB (Schelonka, Graulty, Canseco-Gonzalez, & Pitts, 2017). Here, participants performed a distractor task while unexpected and task irrelevant words and letter symbols were presented within the attended region of space. Through a comparison of ERPs elicited by the unexpected stimuli when they were perceived compared with when they were not perceived, these authors were able to establish which linguistic processes occurred independent of awareness or task relevancy. Results suggested unconscious orthographic processing, with the VAN distinguishing between perceived and not perceived words regardless of the task, and the P3b indexing task-based post-perceptual processing. Critically, Schelonka et al.’s study could not determine whether semantic processing as indexed by the N400 occurred without awareness, as their stimuli (single words) were not designed to elicit an N400, since this requires a stimulus set (words) that includes semantically related and semantically unrelated stimuli. Yet their work provides proof of concept that, with revised methodology, the N400 can be studied during IB.

### The Current Study

The current study sought electrophysiological evidence for semantic processing in the absence of awareness through examining whether the N400 can be elicited during IB. Based on previous electrophysiological work in IB (Pitts et al., 2012; Shafto & Pitts, 2015; Schelonka et al., 2017), we developed a novel paradigm that was designed to maximise the chance to detect an N400 during IB, if one can be observed. We had three concurrent goals. First, we sought to assess whether the N400 can occur unconsciously during IB. Toward this aim, to supplement our univariate ERP analysis, we also used a multivariate decoding procedure to maximise our sensitivity to detect any potential semantic processing differences during IB. Second, we sought to investigate how task relevancy modulates N400 activity. As noted above, previous findings on this issue are mixed (Orgs et al., 2007). Since task relevancy was manipulated across phases two and three of our experiment (the second and third blocks of trials), and because we included measures of stimulus awareness after each phase, we were able to compare the magnitude of N400 activity for identical stimuli that were consciously perceived but were task irrelevant versus task relevant. Third, our IB and task manipulations allowed us to assess the two previously mentioned electrophysiological correlates of consciousness: the VAN and P3b. Previous EEG IB studies have reported the presence of a VAN as a neural correlate of conscious perception of task irrelevant information, whereas P3b is present only once this same information becomes task relevant (Dembski et al., 2021). Because our experimental design used a novel task and stimuli and yet followed the same macro set-up as these previous studies, we were able to examine whether this same pattern of findings occurred within a novel experimental setting.

## Method

### Participants

A total of 53 healthy volunteers (17 male, 36 female; *M* age = 22.79, *SD* = 7.05, range = 18 - 60) participated in the experiment. Nineteen participants were excluded due to either not perceiving any words in both phase one and phase two (*N* = 15), and/or because they did not show an N400 in phase three (*N* = 8) (refer to the Results section for rationale). Of the remaining 34 participants, eight were excluded due to excessive artefacts in their EEG. The final sample for data analysis, 26 participants (5 male, 21 female; *M* age = 21.08, *SD* = 4.02, range = 18 - 30), all reported normal or corrected-to-normal vision and had no history of neurological conditions. All participants provided informed consent prior to participating and were remunerated with $20 AUD. Procedures were reviewed and approved by the university ethics committee (protocol number: 2017/262).

### Stimuli

All stimuli were presented on a white background on a 1920 x 1200 LCD display monitor with a refresh rate of 60 Hz using the Presentation software package (Neurobehavioral Systems). With a viewing distance of approximately 60 cm, participants were presented a central fixation dot within a grid of eight rows of ten pseudo letter characters, each of approx. 0.5° visual angle. A total of 26 pseudo letters were created from Arial Font in Adobe Illustrator via a combination of merging, rotating, or otherwise manipulating letter segments of different characters of the Roman alphabet. The final 26 pseudo letters resembled the Latin alphabet such that real words were camouflaged when presented within the pseudo letter grid, yet were sufficiently dissimilar that they elicited minimal, if any, semantic processing (see Figure 1).

**Figure 1.**
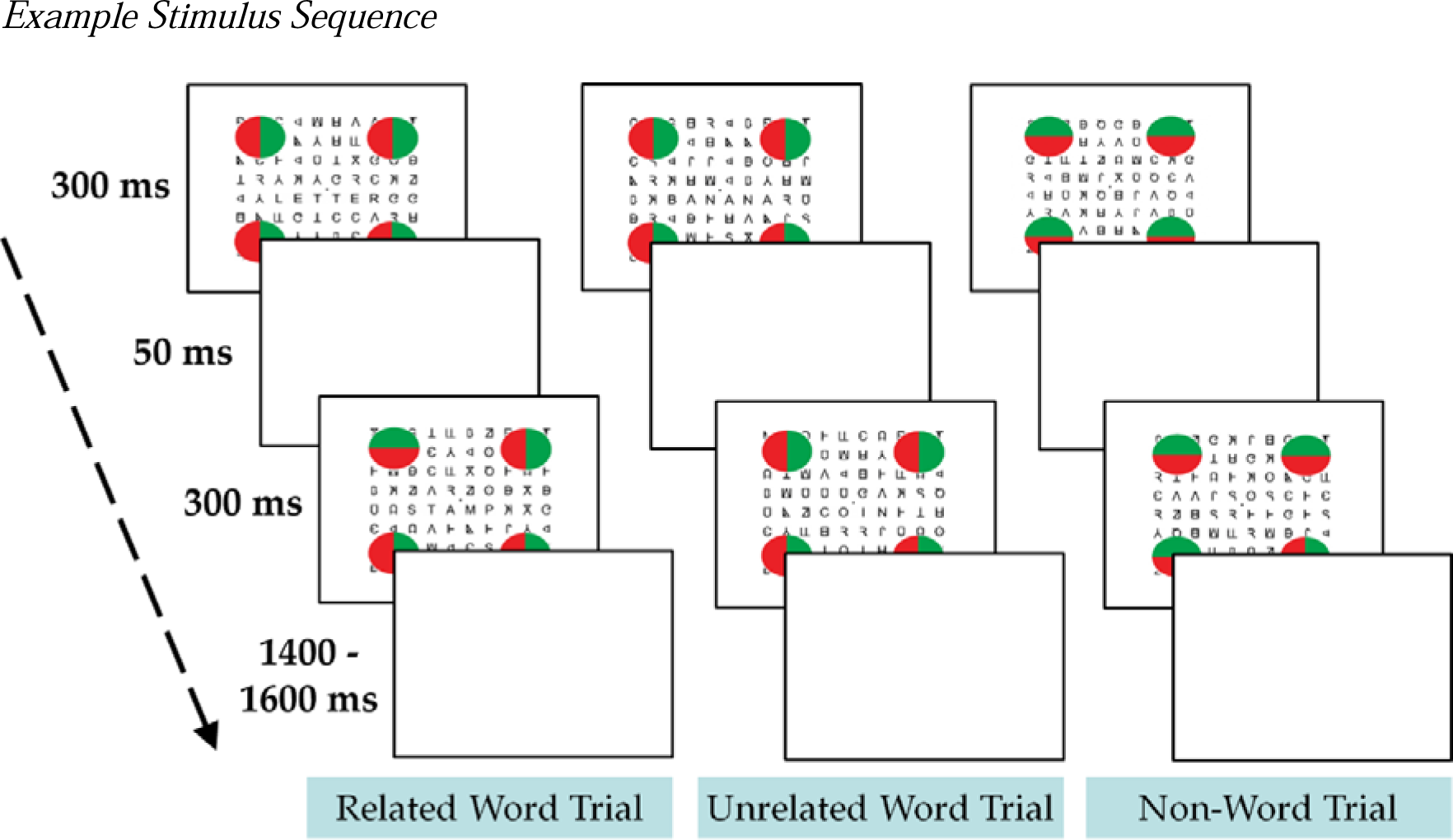
Word pairs were presented centrally, just below fixation. In phase one and two, participants performed a task on the red/green bisected circles (pressing one button if all four circles were the same and another button if one of the circles rotated by 90 degrees). In phase three, participants performed a task on the words (pressing one button for semantically related word pairs, another for semantically unrelated word pairs, and a third button for non-words). Stimuli were identical across the three phases and are shown here for illustration purposes (similar size ratio but not to scale).

A total of 300 words of between four and seven characters in length were chosen based on their length and high frequency of use in American English. Each individual word was paired in two types of sequences: a related pair and an unrelated pair. In the related pair, the target word was preceded by a semantically related prime word (e.g., *letter* followed by *stamp, Figure 1 left*). In the unrelated pair, the target was preceded by a semantically unrelated prime (e.g., *banana* followed by *coin, Figure 1 center*). Each word therefore was presented twice: once as the target in a related pair and once as a target in an unrelated pair, such that our ERP analysis of the N400 (which was time locked to the target) could be compared across related and unrelated pairs by physically identical stimuli.

Semantic relatedness was based on WordNet’s Wu and Palmer (WUP) values, which evaluates relatedness as a product of the proximity of words relative to one another within a set of synonyms (Warin & Volk, 2004). WUP value cut-offs of 0.4 for non-related word pairs and 0.6 for related word pairs were used. A full list of words and their WUP value can be found in the supplementary materials.

Positioned in each corner of the pseudo letter grid were four circles of approx. 1.9° visual angle that were superimposed on top of the pseudo letters. Each circle was bisected such that half was coloured red and half was coloured green. During the first stimulus presentation within a given sequence, all four circles were identically oriented (i.e., the bisection of all circles was either vertical or horizontal). During the second stimulus presentation within a given sequence, the orientation of the circles either remained unaltered or one of the circles were rotated by 90 degrees such that its bisection was perpendicular to the other discs (see Figure 1 for examples).

A single stimulus sequence lasted between 2,050 to 2,250 milliseconds: two 300 millisecond pseudo letter grid presentations interspersed by a 50-millisecond blank screen, followed by a variable 1,400 to 1,600 millisecond blank response window. There were three types of stimulus sequences: a related word sequence, an unrelated word sequence, and a non-word sequence (see Figure 1). In the former two, the centre-most grid positions immediately below fixation contained word stimuli. Each block of trials always began with a non-word sequence, and word sequences were interspersed between one or two non-word sequences such that there were approx. 1.5 times as many non-word sequences as there were word sequences in any given trial. A block of trials lasted approx. 1.7 to 1.9 minutes and consisted of approximately 50 stimulus sequences: ten related word sequences, ten unrelated word sequences, and ∼ 30 non-word sequences. There was a total of ten blocks of trials per phase, and three phases in total. Stimuli were identical in all phases of the experiment but presented in a different random order within each phase.

### Procedure

The experiment consisted of three phases that involved two separate tasks: a change detection task involving the bisected circles in phase one and phase two, and a semantic judgment task involving the word pairs in phase three. This shift in task enabled the critical manipulation of task relevancy: words were task relevant in phase three but remained task irrelevant in phase one and phase two. For the change detection task, participants responded with a left arrow button response if no circles had rotated and a right arrow button response if any one of the circles had rotated. For the semantic judgment task, participants were instructed to ignore the colored circles and respond whether, within a single sequence, a related word pair (left arrow button response), unrelated word pair (down arrow button response), or non-word pair (right arrow button response) was presented. Participants were given self-paced rest breaks after every block of trials. Each phase lasted between approx. 17 and 25 minutes.

Prior to the experiment, participants were provided examples of what to look for in the colored circle task. Participants then ran through five practice trials, during which no word pairs were presented. Following phase one, participants were provided a questionnaire in which they were asked to detail whether they had noticed anything unexpected during the experiment. The questionnaire served to both determine whether participants had spontaneously noticed the words and to cue participants that words would appear within the pseudo letter grid in subsequent phases of the experiment. The questionnaire presented participants with example items that included both real (e.g., a word) and foil stimuli (random string of consonants, random string of numbers, several coloured squares). Using a five-point scale, participants rated both their confidence (1. *very confident I did not see it*, 2. *confident I did not see it*, 3. *uncertain*, 4. *confident I saw it*, 5. *very confident I saw it*) and frequency (1. *never*, 2. *rarely, e.g. less than 10 times*, 3. *infrequently, e.g. 10-50 times*, 4. *frequently, e.g. 50-100 times*, 5. *very frequently, e.g. over 100 times*) of having seen the item in the previous phase. Participants were considered inattentionally blind if they rated their confidence in seeing words as a three or less, and/or if they provided no evidence of noticing any words (e.g., during the open-ended response). Participants received the same questionnaire following phase two.

### EEG Pre-Processing and Analysis

Data were recorded at a sample rate of 2056 Hz using a 64-channel BioSemi Active Two system. As is standard for the BioSemi system, the Common Mode Sense and Driven Right Leg served as ground during recording. External electrodes were applied to the left and right mastoids, the outer canthi of each eye, and above and below the left eye. Eye electrodes were re-referenced offline to single horizontal and vertical eye channels.

All pre-processing was completed using custom Matlab scripts written using EEGlab (Delorme & Makeig, 2004). Raw data and all analysis scripts are available at: https://osf.io/gpxub/. Raw data were first down sampled to 256 Hz and re-referenced to the average of the left and right mastoids. A notch filter was then used to remove 50 Hz line noise. Noisy channels were identified based on a kurtosis value exceeding 5 standard deviations of the norm and were then interpolated using spherical spline interpolation. Next, data were band-pass filtered from 0.1 to 30 Hz using a fourth order, zero-phase Butterworth filter. For analysis of each ERP, data were epoched into 1200 millisecond time windows, from 200 milliseconds pre-stimulus to 1000 milliseconds post-stimulus. Investigation of the N400 and P3b involved analysis time-locked to the second stimulus (words only for the N400; words and non-words for the P3b), while for the VAN, analysis were time-locked to the first stimulus (words and non-words). We elected to examine the second stimulus (the target) for the P3b, rather than the first stimulus (the prime), since the presentation of the target overlapped the time-course of the P3b to the first stimulus. ERPs were baseline corrected by subtracting mean voltages within the pre-stimulus to stimulus onset interval from every data point. Trials containing artefacts (e.g., blinks, eye movements, muscular noise) were detected and removed using the ERPlab functions *artmwppth* (for detection of blinks), *artstep* (for detection of eye movements), and *artextval* (for all other artefacts) using amplitude thresholds of 100 µV, 50 µV, and +/− 75 µV, respectively (with adjustments to these thresholds as needed at the participant level). Participants were excluded if they had less than 30 trials per condition for the primary N400 analysis.

For ERP analysis, electrode channels for each component (N400, P3b, VAN) were selected based on previous literature (Šoškić et al., 2022). A group of central channels were used for the N400 (Cz, CPz, C1, CP1, C2, CP2), parietal channels for the P3b (Pz, P1, P3, P5, P7, P9, P2, P4, P6, P8, P10), and parieto-occipital channels for the VAN (O1, O2, PO3, PO7, PO4, PO8). The N400 component was measured based on the difference between related and unrelated target words, while the P3b and VAN were measured based on the difference between words and non-words (targets for the P3b and primes for the VAN). Mean amplitude over a 500ms time window (500ms to 1000ms) was selected for analysis of the N400 and P3b, while a 100ms time window was selected for the VAN (320ms to 420ms).

All decoding was performed using the MVPA light toolbox in Matlab (Treder, 2020). Epoched data were first low pass filtered at 6 Hz and resampled at 50 Hz. All external channels were removed (such that all 64 scalp channels remained) and datasets were then reformatted from a channels by time by trial structure, to a trials by channel by time structure, and labelled according to whether the trial corresponded to a related or unrelated word pair. As part of MVPA light’s built-in preprocessing parameters, we applied *z-scoring* (demeaning and rescaling of the data), *average sampling* (increasing the signal-to-noise ratio via splitting samples from the same class into multiple groups and then replacing each group with the mean of that group), and *under sampling* (in the instance that there is an imbalance in the samples between conditions, samples are removed to equate the number of samples between conditions). We used default parameters for each. For average sampling, this meant that 5 trials went into each pseudo-average. For our classifier, we used linear discriminant analysis (LDA) with five iterations of a four-fold cross validation procedure. We implemented LDA toward both decoding of the time series (where the classifier is trained on each time point and tested on the same time point) and temporal generalisation (where the classifier is trained on every time point, and then each time point is tested based on training from every other time point). We assessed model performance with area under the curve (AUC), which is similar to accuracy but takes into account false alarms and misses. For statistical analyses, decoding performance was assessed using a Wilcoxon rank test performed at every time point, with a cluster-based permutation for correction for multiple comparisons (1,000 permutations).

### Statistical Analysis

Statistical analyses on the ERPs were conducted using R version 4.2.0 (R Core Team, 2021) and JASP version 0.14.1 (JASP team, 2022). Both frequentist and Bayesian statistics are reported for all analyses (Keysers et al., 2020). For frequentist statistics, we applied the standard threshold of *p* < 0.05 (two-tailed). Effect sizes are reported as Cohen’s *d* or partial-eta squared (η_p_^2^). For Bayes statistics, Bayes factor (BF) was computed using Rouder’s method with the default Cauchy prior of 0.707 for *t*-tests, *r* = 0.5 for fixed effects, and *r* = 1 for random effects in ANOVAs (Morey & Rouder, 2011; Rouder et al., 2012). We report BF_10_, which indicates the likelihood of the alternative hypothesis compared with the null hypothesis. Evidence for the alternative hypothesis was interpreted as inconclusive for BF_10_ < 3, moderate for BF_10_ ≥ 3, strong for BF_10_ ≥ 10, very strong for BF_10_ ≥ 30, and extreme for BF_10_ ≥ 100 (Jeffreys, 1961). For analysis of behavioural data (accuracy, reaction time), outliers were identified and removed using Tukey’s method (1977).

## Results

Of the final 26 participants, all reported being unaware of the words in phase one, whereas all participants reported noticing the words in phase two (see Figure 2). One participant gave a confidence rating of three in phase two but was classified as aware as their frequency rating and open-ended response indicated that they had seen the word stimuli.

**Figure 2.**
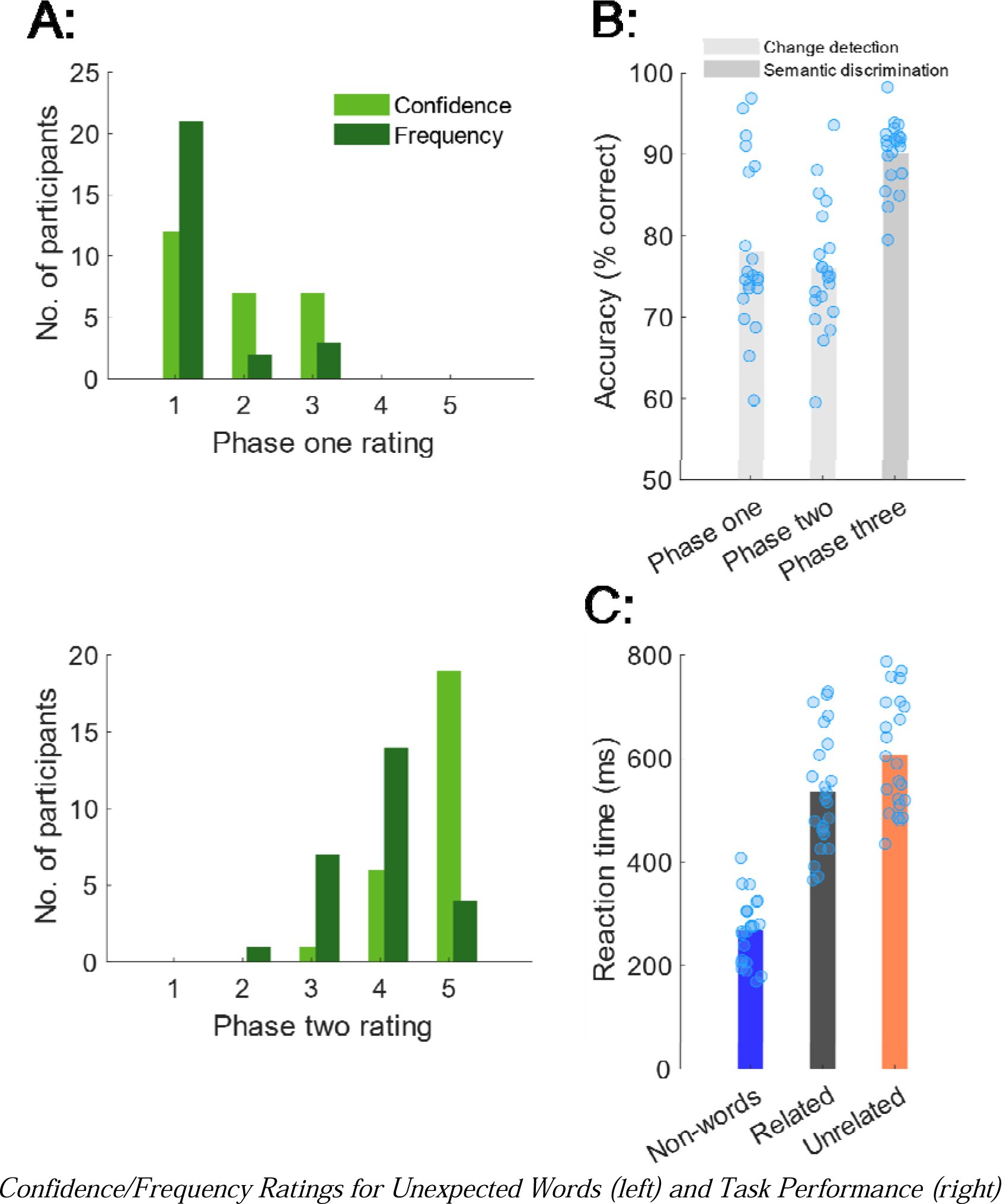
**A)** Confidence/frequency ratings from the post-phase questionnaire on noticing the unexpected stimuli (words), where 1 on the x axes reflects the lowest confidence/frequency rating and 5 reflects the highest confidence/frequency rating. The top chart shows post-phase questionnaire data following phase one and the bottom chart shows post-phase questionnaire data following phase two. **B)** Accuracy across phase one (change detection task), phase two (change detection task), and phase three (semantic discrimination task). **C)** Reaction times during phase three, when participants performed the semantic discrimination task and had to respond based on whether the stimulus pair were non-words, a related pair of words, or an unrelated pair of words. For panels **B** and **C**, the blue scatter shows individual participant data points.

### Behavioural Performance

*Accuracy.* A repeated measure ANOVA examining behavioural accuracy across phases (phase one, phase two, phase three) revealed a significant effect for accuracy, *F*(2,40) = 22.94, *p* < .001, η_p_^2^= .53, BF_10_ > 100. Follow up analyses confirmed that participants performed better at the semantic judgment in phase three than the change detection task in phase one, *t*(20) = 5.26, *p* < .001, *d* = 1.15, BF_10_ > 100, and phase two, *t*(20) = 6.71, *p* < .001, *d* = 1.46, BF_10_ > 100. The difference between task performance in phase one and phase two was not significant, *t*(20) = 0.90, *p* = .38, *d* = 0.20, BF_10_ = 0.33 (see Figure 2B).

*Reaction Time.* Separate repeated measures ANOVAs were performed for each phase to examine differences in reaction time based on word condition (related, unrelated, non-words). The main effect of word condition was not significant in phase one, *F*(2, 44) = 2.76, *p* = .07, η_p_^2^ = .11, BF_10_ = 0.90, or phase two, *F*(1.46, 32.06) = 0.42, *p* = .60, η_p_^2^ = .02, BF_10_ = 0.17. There was a significant effect for word condition in phase three, *F*(1.23, 27.06) = 172.56, *p* < .001, η_p_^2^ = .89, BF_10_ > 100. Follow up analyses confirmed that in phase three, participants responded faster to non-words than to related words, *t*(22) = 11.83, *p* < .001, *d* = 2.47, BF_10_ > 100, and to unrelated words, *t*(22) = 14.86, *p* < .001, *d* = 3.10, BF_10_ > 100.

Related words were also responded to faster than unrelated words, *t*(22) = 8.05, *p* < .001, *d* = 1.68, BF_10_ > 100 (see Figure 2C).

### Is the N400 Elicited Under Conditions of Inattentional Blindness?

For analysis of the N400, the primary contrast involved comparing ERPs time-locked to the second word of related word pairs compared with unrelated word pairs. This same contrast was made separately for each of the three phases of the experiment. We expected a robust N400 in phase three, since stimuli in that phase were consciously perceived and task relevant. Phase three therefore served as a validation check to ensure our design could reliably elicit an N400. A significant (even if reduced in magnitude) N400 in phase two would indicate semantic processing of minimally attended, consciously perceived, but task irrelevant words. A significant N400 in phase one would imply semantic processing of stimuli that were fully unattended and not consciously perceived (since no participants in this phase reported noticing the word pairs).

A 2 (relatedness: related, unrelated) by 3 (phase: phase 1, phase 2, phase 3) repeated measures ANOVA comparing mean amplitudes (500-1000ms) over central electrodes indicated a main effect for relatedness, *F*(1,25) = 25.62, *p* < .001, η_p_^2^ = .51, BF_10_ > 100, a main effect for phase, *F*(2, 50) = 30.55, *p* < .001, η_p_^2^ = .55, BF_10_ > 100, and an interaction between relatedness and phase, *F*(2,50) = 20.43, *p* < .001, η_p_^2^ = .45, BF_10_ > 100. Follow up analyses comparing related with unrelated words for each phase separately suggested significant effects in phase three, *t*(25) = 6.93, *p* < .001, *d* = 1.28, BF_10_ > 100, and phase two, *t*(25) = 2.66, *p* = .01, *d* = 0.52, BF_10_ = 3.71, but not in phase one, *t*(25) = 0.28, *p* = .78, *d* = 0.06, BF_10_ = 0.21. Waveforms and scalp maps summarizing these results are provided in Figure 3.

**Figure 3.**
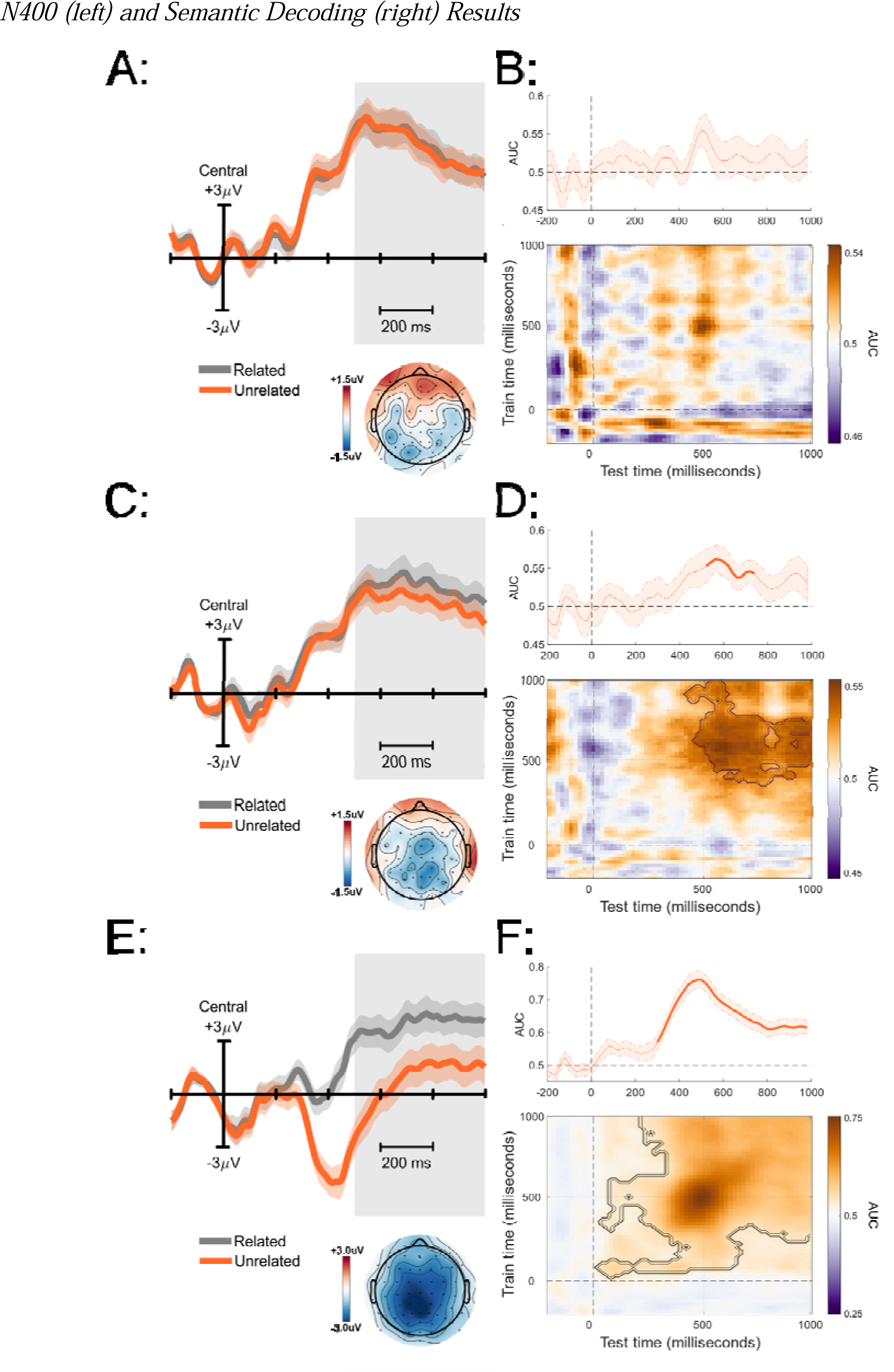
Grand-averaged event related potentials, scalp topographies, and decoding performance for the related and unrelated word conditions for phase one **(A-B)**, phase two **(C-D)**, and phase three **(E-F)**. Time zero in all plots reflects the onset of the second word in each pair. ERPs are pooled over the central electrodes (Cz, C1, C2, CPz, CP1, CP2) used to quantify the N400 wave. The shaded area in the waveform plots **(A, C, E)** indicate the time window used for statistical analysis. Scalp maps show the difference in voltage topography averaged over the analysis time window, calculated by subtracting the unrelated condition amplitudes from the related condition amplitudes. Shaded regions around the ERP waveforms and decoding curves indicates +/− 1 standard error of the mean. Decoding results **(B, D, E)** are separated into time series decoding (top of **B,D,E**) and temporal generalisation matrixes (bottom of **B,D,E**). Performance reflects average decoding accuracy at each time point. Bolded regions on the time series plots (top) and regions outlined in the temporal generalisation matrixes (bottom) indicate timepoints in which decoding accuracy was significantly above chance (corrected for multiple comparisons).

Next, to assess the magnitude of N400 modulation across phases, we ran a repeated measures ANOVA comparing difference-related activity (unrelated minus related amplitudes) across phases of the task (phase one, phase two, phase three). Results indicated a significant effect for phase on N400 amplitudes, *F*(2,50) = 20.43, *p* < .001, η_p_^2^ = .45, BF_10_ > 100. This was due to differences between phase one and phase three, *t*(25) = 5.99, *p* < .001, *d* = 1.18, BF_10_ > 100, and phase two and phase three, *t*(25) = 4.34, *p* < .001, *d* = 0.85, BF_10_ > 100 (see Figure 3D).

Overall, there were clear differences in the processing of related and unrelated words, as reflected by the presence of an N400, when participants were aware of the stimuli in phases two and three. Task relevancy of the word pairs modulated the amplitude of the N400, but this signal was present regardless of the task. However, the N400 was absent when participants were inattentionally blind to the word pairs in phase one, and Bayes factors suggested that there was moderate evidence for a null effect of semantic relatedness in this phase (BF_10_ = 4.76). Our ERP results therefore suggest that no semantic processing occurred in the absence of awareness due to inattention. Mean amplitudes for N400 activity are presented in Table 1.

**Table 1:** Mean amplitude and standard deviation for ERP components.

|  | N400 | VAN | P3b |
| --- | --- | --- | --- |
| Phase 1 | -0.11 (1.95) | -0.17 (0.75) | 0.14 (0.84) |
| Phase 2 | -0.80 (1.53) | -0.71 (1.08) | -0.08 (1.23) |
| Phase 3 | -3.18 (2.48) | -0.30 (1.92) | 2.00 (2.82) |

Note. Values indicate amplitudes (uV) of difference-related activity for each ERP component for each phase of the task. Standard deviations are in parentheses.

### Is Semantic Relatedness of Word Pairs Decodable During Inattentional Blindness?

Neural decoding results for each phase are provided in Figure 3. In phase three, the semantic relatedness of the word pairs was decodable within the time series data from 300ms post stimulus and peaked with an accuracy of 76% at ∼ 500ms (*p* < .001). Despite a reduction in accuracy from 76% to 62% from 500ms to 820ms, decoding performance remained significantly above chance until the end of the epoch. When the decoder was trained and tested time point by time point, strong temporal generalisation was observed that spanned nearly the entire epoch, with an onset that preceded the above chance time series decoding by about 259ms, onsetting at ∼41ms and lasting until the end of the epoch (*p* < .001). In phase two, decoding accuracy for the time series reached significantly above-chance levels from 560ms to 740ms (*p* = .01), with a peak in accuracy (55%) at 680ms. When the decoder was trained and tested time point by time point, some temporal generalisation was apparent in these later time periods where there was a significant burst of activity (*p* = .02). Despite promising decoding performance in both phase two and phase three, semantic relatedness was not decodable at any timepoint in phase one (see Figure 3A). Results from multivariate decoding were therefore consistent with the univariate ERP measures: significant effects were present within the N400 timeframe in phase two and phase three, but not phase one. In combination with our ERP results, these data provide evidence that information regarding the semantic relatedness of word pairs was not present in the EEG signal during phase one when participants were not aware of the words.

### Are Electrophysiological Signatures of Conscious Perception Present in this Modified Inattentional Blindness Paradigm?

To test for the presence of the “visual awareness negativity” (VAN) in the task irrelevant conditions, a 2 (word condition: word, non-word) by 2 (phase: phase 1, phase 2) repeated measures ANOVA with mean amplitudes (320-420ms) over parieto-occipital areas was conducted. This analysis revealed a significant main effect for word condition, *F*(1, 25) = 5.55, *p* = .03, η_p_^2^ = .18, BF = 5.69, and a significant main effect for phase, *F*(1.41, 35.23) = 8.07, *p* = .003, η_p_^2^= .24, BF_10_ = 0.62. Follow-up comparisons comparing words with non-words for each phase separately indicated a significant difference between words and non-words in phase two, *t*(25) = 3.36, *p* = .002, *d* = 0.66, BF_10_ = 15.48, but not phase one, *t*(25) = 1.18, *p* = .25, *d* = .23, BF_10_ = 0.38. Further analysis confirmed that there was a significant difference in difference-related activity (words minus non-words) between phase one and phase two, *t*(25) = 2.34, *p* = .03, *d* = 0.46, BF_10_ = 2.04, corresponding with an effect of awareness across phases of the task. These results are summarized in Figure 4.

**Figure 4.**
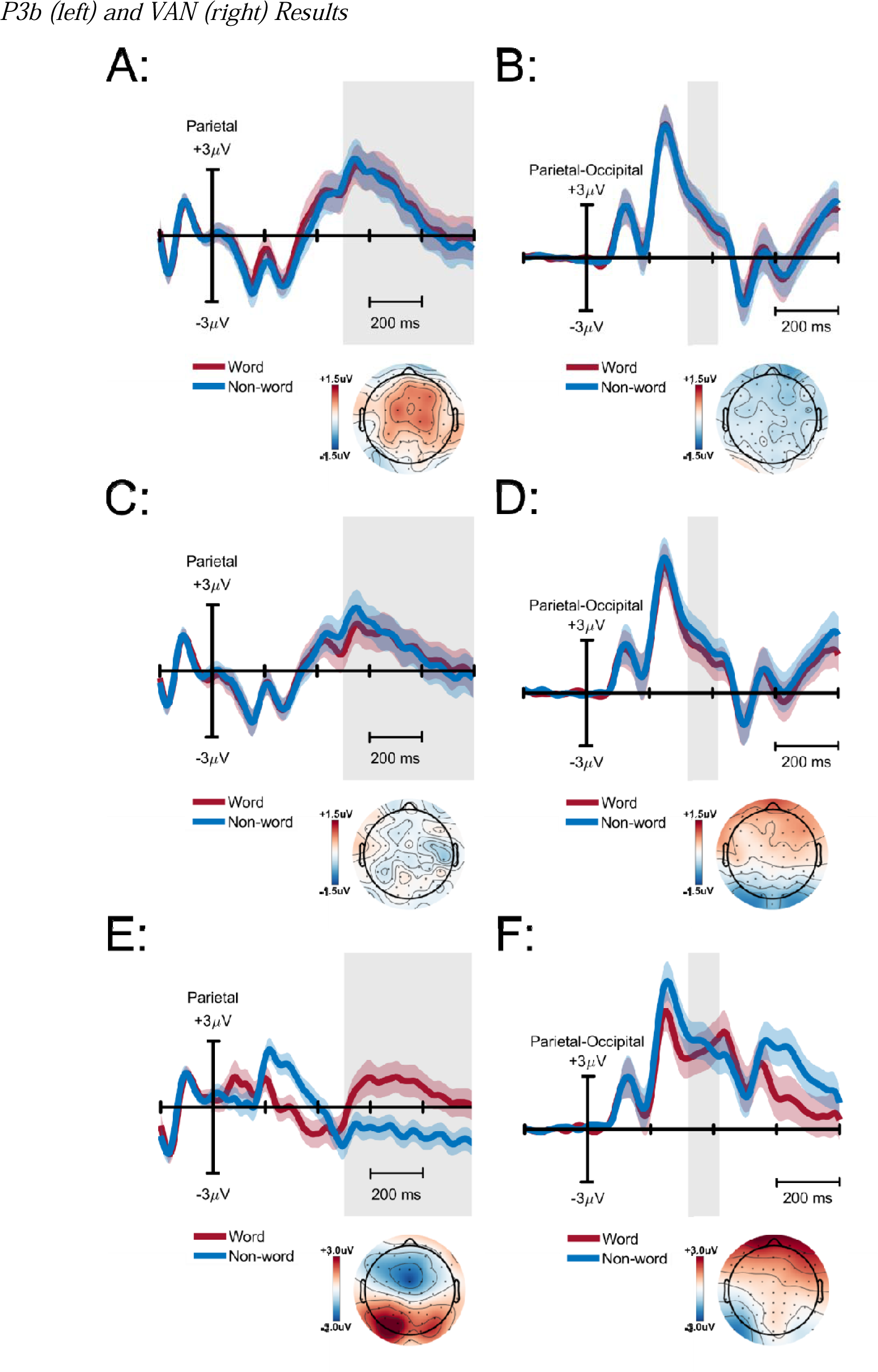
Grand-averaged event related potentials and scalp topographies for the word and non-word conditions for phase one **(A-B)**, phase two **(C-D)**, and phase three **(E-F)**. Plots on the left **(A, C, E)** show P3b results and are pooled over the parietal electrodes (Pz, P1, P3, P5, P7, P9, P2, P4, P6, P8, P10) used to quantify the P3b. Time zero in these plots is the onset of the second (non-)word in each pair. Plots on the right **(B, D, F)** show VAN results and are pooled over the parietal-occipital electrodes (O1, O2, PO3, PO4, PO7, PO8) used to quantify the VAN. Time zero in these plots is the onset of the first (non-)word in each pair. The shaded area in all waveform plots indicate the time window used for statistical analysis. Scalp maps show the difference in voltage topography averaged over the analysis time window, calculated by subtracting the non-word condition amplitudes from the word condition amplitudes. Shaded regions around the ERP waveforms indicate +/− 1 standard error of the mean.

To assess the P3b, a 2 (word condition: word, non-word) by 3 (phase: phase 1, phase 2, phase 3) repeated measures ANOVA with mean amplitudes (500-1000ms) over parietal regions was conducted. This analysis indicated a significant main effect for condition, *F*(1, 25) = 7.00, *p* < .001, η_p_^2^ = .22, BF_10_ = 2.29, a main effect for phase, *F*(1.31, 32.76) = 6.75, *p* = .009, η_p_^2^ = .21, BF_10_ = 17.12, and an interaction between word condition and phase, *F*(1.33, 33.27) = 13.78, *p* < .001, η_p_^2^ = .36, BF_10_ > 100. Follow-up tests comparing words with non-words in each phase separately indicated a significant difference between words and non-words in phase three only, *t*(25) = 3.61, *p* = .001, *d* = .71, BF_10_ = 26.88. The P3b was not present in phase two, *t*(25) = 0.34, *p* = .74, *d* = .07, BF_10_ = 0.22, or phase one, *t*(25) = 0.85, *p* = .40, *d* = .17, BF_10_ = 0.29. Figure 4 summarizes these results and mean amplitudes for the VAN and P3b are presented in Table 1.

In sum, these results corroborate previous findings from other EEG-adapted IB studies that VAN reflects an early electrophysiological signature of conscious perception, even for task-irrelevant stimuli (i.e., phase two versus phase one of the current experiment), whereas the P3b is linked with consciously perceived stimuli that are task relevant (i.e., in phase three of the current experiment).

## Discussion

The current study sought electrophysiological evidence for semantic processing of stimuli that were not consciously perceived due to inattentional blindness. We presented participants with pairs of words that were either semantically related or unrelated while they simultaneously performed an attentionally-demanding task involving non-word stimuli. As participants’ were distracted by the primary task and were not informed of the presence of the words prior to the experiment, we were able to present the words many times whilst they remained unseen. Our results demonstrate that electrophysiological signatures of semantic processing were not observable in phase one of the task, when all participants were unaware of the words due to inattentional blindness. In phase two, when the word pairs were partially attended and consciously perceived, but remained task irrelevant, we found both an N400-effect and above-chance decoding of words based on their relatedness. In phase three, when the same words were fully attended and task relevant, a large N400 was observed and decoding performance was considerably enhanced. These findings align with some previous literature that has reported an absence of the N400 when participants do not consciously perceive the stimuli (Batterink et al., 2009; Holcomb & Grainger, 2009; Kang et al., 2011) and conflicts with other work that suggests semantic processing might still occur for stimuli that are not perceived due to inattention (Schnuerch et al., 2016; Pugnaghi et al., 2020).

### No Evidence for Unconscious Semantic Processing?

Is it possible that unconscious semantic processing is, in fact, detectable in scalp-based EEG measures, but such effects were too weak or variable in our paradigm? We find this unlikely for three reasons. First, Bayes factor in phase one suggested moderately strong evidence for the null (BF_10_ = 4.76), that is that there was no difference in ERP amplitudes between related and unrelated word pairs (Jarosz & Wiley, 2014). Second, by design, the stimuli and task in phase two were identical to phase one. The only difference between these phases was the small change in attention and expectations cued by the intervening awareness questionnaire. Thus, the fact that an N400 was present in phase two confirms that such a signal is detectable with the current methods of analyses in this paradigm. Third, if the semantic effect in phase one was too variable in time or scalp distribution across participants to be present within standard univariate statistical analyses, these differences may still be detectable when employing more sensitive multivariate analyses, such as EEG decoding (Bae & Luck, 2018). Despite the improved sensitivity of EEG decoding for uncovering informational differences across the whole scalp, our decoding results were consistent with and corroborated our N400 results. We found significantly above chance decoding in both the time series and temporal generalisation matrices for phase two and phase three—precisely where significant ERP differences were observed—but not phase one, where ERP differences were minimal. Therefore, both our ERP and decoding results converge on the conclusion that information regarding semantic processing was absent from participants’ EEG signal when they were not aware of the eliciting stimuli due to inattention. Though it should be noted that it remains possible that unconscious semantic processing can occur, but the relevant neural signals are only measurable using more precise neural recording techniques, such as intracranial EEG or single-unit recordings.

Relatedly, it might be that the absence of an N400 in phase one of our experiment was due to participants’ perceiving a single of the words in each pair (but not both). In previous studies, the N400 has been found if the target (second) word is masked or unattended and the prime word is clearly perceived (Holcomb & Grainger, 2009; Stenberg et al., 2000). For this reason, some have recommended employing designs that allow for explicitly testing whether the prime was perceived when studying the N400 (Holcomb & Grainger, 2009). However, because participants were only able to be questioned on their perception of the words after an entire phase was completed, we were unable to gauge participants’ awareness of the words on a trial-by-trial basis; and by extension, whether the first (the prime) or second (the target) word was perceived in any given stimulus sequence. Yet, this issue would only be of concern if we had found no N400 in phase two, since it is under these circumstances that the absence of an N400 might be due to participants having perceived only one of the words (either the prime or target). Since our null findings occurred in phase one, where all participants reported having *not* perceived any words at all, this explanation does not hold weight here. Moreover, any differential awareness of the prime compared with the target in the subsequent phases of the task (when all participants perceived the words) would, if anything, *reduce* the size of the N400 (if indeed such a confound would affect N400 magnitude in the first place). We can confidently state then that this was not a problem in these experimental phases, and our study more broadly, since we observed an N400 in both. In sum, it seems unlikely that any differential awareness of individual words impacted our findings.

A more pressing concern arises when reflecting upon how our findings fit with recent work that reported behavioural evidence of semantic processing during IB. Schnuerch et al. (2016) and Pugnaghi et al. (2020) found that participants’ performance in a numerical categorisation task was hindered when unexpected number stimuli were simultaneously presented that were incongruent, compared with when they were congruent, with the category of the target number. Despite that participants were inattentionally blind to the unexpected numbers, some degree of semantic categorization of the unexpected critical stimuli must have occurred for performance on the primary task to have been hindered. Though these findings are at odds with our own, it is worth noting that semantic-related reaction time effects have previously been shown to persist for unconscious stimuli, even when the N400 did not (Brown & Hagoort, 1993).

Besides outcome measures, a critical difference between studies that may have contributed to differences in results lay in the attentional control settings of participants. In the current study, word pairs were – by design – irrelevant to the task participants were initially instructed on, and hence were not congruent with what attention was “set” to (Hutchinson et al., 2022). In contrast, in Schnuerch et al. (2016) and Pugnaghi et al. (2020), for any effect to have occurred, findings necessitated that the unexpected stimuli were congruent with what participants’ attention was set to. This discrepancy may explain the difference in results obtained between studies. Given that participants were explicitly required to respond to numbers, the critical stimuli in Schnuerch et al. (2016) and Pugnaghi et al. (2020) were likely apportioned some degree of implicit task relevancy, which may have facilitated unconscious semantic processing through pre-conscious attentional allocation. No such benefit would have occurred for word stimuli in the current study. This interpretation clarifies differences in study findings and reemphasizes the need to account for task relevancy when probing the N400. Perhaps it is possible that semantic processing can occur without awareness of the eliciting stimuli, however our results suggest this is not the case for stimuli that are not perceived due to inattention and which are concurrently task irrelevant.

Follow up work could therefore look to establish whether any tentative unconscious semantic processing effect varies, as we predict, when attention set is manipulated. A future variation of the current experiment could, for example, involve letter or word stimuli as part of the primary task in the first two phases, instead of bisected colour circles. Conversely, we would predict the interference effect observed in Schnuerch et al. (2016) and Pugnaghi et al. (2020) to disappear if the critical stimuli were varied to be incongruent with the observers’ attention set.^1^

A critical point need be noted that the precise mechanism underlying how attention set moderates IB is, at present, not entirely clear (Hutchinson et al., 2022). Stimuli that are incongruent with an observer’s attention set may be susceptible to IB simply due to an absence of any pre-conscious attentional allocation. Alternatively, stimuli may instead be actively suppressed from being perceived. This distinction is important in the current context, in consideration of work that has found that task instructions can attenuate the N400. In Erlbeck et al. (2014), participants were to either explicitly attend to, passively ignore, or actively ignore auditory sentences. Their experimental conditions were conceptually similar to our own: in the explicitly attend condition, stimuli of which the N400 was measured were task relevant, as per phase three in the present study. Conversely, in the active ignore condition, participants attended to an alternative task and hence the eliciting stimuli were task irrelevant—as per phase one and phase two in the current study. Notably, the N400 was markedly smaller in the passive ignore condition and was entirely absent in the active ignore condition (Erlbeck et al., 2014). An unconscious semantic processing effect may therefore have persisted in Schnuerch et al. (2016) and Pugnaghi et al. (2020) because their stimuli were not susceptible to active suppression (due to being congruent with the observer’s attention set). Stimuli in the present study, on the other hand, were incongruent with what observers were attending to and consequently may have been vulnerable to active suppression, which may have attenuated any N400 activity (Erlbeck et al., 2014).

That high rates of IB were observed in the work of Schnuerch et al. (2016) and Pugnaghi et al. (2020), despite an attention set match between the unexpected stimuli and task relevant items, implies that IB was induced in these studies through some combination of task difficulty and perceptual load.^2^ That an effect persisted in their work is therefore quite remarkable, however neither task difficulty nor perceptual load have been found to impact semantic processing—at least with respect to the N400 (Kemp et al., 2019; Neumann & Schweinberger, 2009). Though it is worth noting that Pugnaghi et al. manipulated perceptual load, finding that an interference effect remained under conditions of high load when the unexpected stimuli were incongruent with the primary task. Critically, there was no interaction between load and congruency. This suggests that excessive load *did* abolish semantic processing, as unconscious semantic processing would require that the largest reduction in task performance occurred in the high load semantic incongruent condition. That this was not observed suggests excessive load interfered with semantic processing but left some low-level processing of the unexpected stimuli intact. Evidently, task load can interfere with unconscious semantic processing, and hence there may still be scope to evaluate its role in future work on the N400 under conditions of IB.

### Task Irrelevant Semantic Processing

The presence of an N400 in phase two of the current study was not a foregone conclusion. While all participants in this phase of the experiment self-reported noticing the word stimuli, they were nevertheless still performing the distracting change detection task on the circular discs. Hence, the presence/absence of an N400 in this phase was somewhat contingent on the broader question of whether the N400 reflects controlled or automatic processes (Kutas & Federmeier, 2011). This is not entirely analogous to the question of whether the N400 can occur without awareness, since the automaticity of N400 is typically thought to hinge on attention—automatic processes are those that can occur even without effortful endogenous attention directed to the stimulus, while controlled processes require some degree of explicit goal-driven attention. The fact that an N400-effect was observed in phase two, when word pairs were not afforded any such goal-driven attention, therefore speaks to this issue, as it suggests at least some automaticity to the generation of N400, but with the requirement of at least some minimal amount of spatial (or object-based) attention. We assume such attention was cued by the questioning in between phases and is what resulted in participants becoming aware of the words in phase two.

We were also able to estimate an effect size for the effect of task relevancy on the N400 since any difference in amplitude between phase two and phase three would be driven by the change in task relevancy of word stimuli. Note that differences in stimuli themselves could not explain differences across phases, since the same word pairs were used for all three phases of the experiment. If anything, our effect size estimates for across phase effects is thus an underestimate, since stimulus repetition attenuates N400 magnitude (Kutas & Federmeier, 2011). The effect size we estimated (*d* = 0.85) suggests that task relevancy has a large effect on the N400, yet because a task irrelevant N400 was still observed, task irrelevancy is not sufficient to abolish the N400 altogether.

Several previous works have examined N400 activity for task irrelevant stimuli. In early work by McCarthy and Nobre (1993), the degree to which attention modulated the N400 could be compared by varying where word pairs were presented relative to the loci of spatial attention. Similar to current work, participants were informed about the presence of task relevant items requiring a response, and hence unattended stimuli that failed to elicit an N400 were task irrelevant. However, they did not report whether participants perceived the task irrelevant words, therefore it is unclear whether the N400 was attenuated due to task irrelevancy or because these stimuli were not perceived. In the current study, we were able to clearly differentiate between the effect of awareness and task relevancy on the N400.

Moreover, unlike McCarthy and Nobre (1993), words in the current study were presented within the spatially attended region, directly below where participants were foveating therefore limiting any possible contamination of effects due to spatial attention constraints. Later work by Relander et al. (2008) found significant N400 activity for task irrelevant spoken words, yet similar to the current study, waveforms elicited by task irrelevant stimuli were visually dissimilar from those elicited when stimuli were task relevant. In the current study, these differences (e.g., the absence of the classic N4 peak in phase two) likely resulted from the fact that participants were performing a separate task, therefore any semantic processing of the word pairs was considerably delayed and/or not prioritised in this phase.

It is worth noting that some of the N400 effects for task irrelevant stimuli found in previous studies may occur due to either task switching across trials or order effects. Semantic processing may occur during trials in which the eliciting stimuli are task irrelevant due to carry over effects from the previous trial or prior knowledge of their relevancy (Orgs et al., 2007, 2008). These same criticisms do not hold in our study since the task switch occurred only in phase three (after two experimental phases) and the change detection task always preceded the relatedness judgment task. As such, there was no possible way for participants to allot any form of task relevancy to the words in either phase one or phase two, as participants did not know at these stages that words would later require a response. In sum, then, while our data suggest the N400 requires awareness, it nonetheless seems to be elicited through some degree of automatic processing that can occur even when the stimuli are irrelevant to the current goals of the observer, at least within the visual modality.

### Electrophysiological Correlates of Consciousness

We observed electrophysiological correlates of consciousness in corroboration with prior works in similar paradigms (Pitts et al., 2012; Schnuerch et al., 2017; Harris et al., 2018). In phase two, when all participants were aware of the words, we observed a subtle yet distinct difference occurring ∼320ms post stimulus over parietal-temporal-occipital regions of the scalp. This same difference was not apparent in phase one, when all participants were unaware of word pairs. This difference is commonly referred to as the “visual awareness negativity” (VAN) and has been previously found in phase two of similar IB tasks, in addition to the aware group of participants in phase one (which we were not able to test for here as all participants were unaware of the words in this phase) (Shafto & Pitts, 2015). By comparison, P3b only occurred in phase three, when these same stimuli were perceived and task relevant.

Each ERP has been considered as evidence for / against competing theories of consciousness. For example, some consider the VAN as evidence for so-called “sensory” theories of consciousness, while the P3b is taken as evidence for the global neuronal workspace model (Dehaene & Naccache, 2001; Forster et al., 2020). Our findings suggest the VAN is a better candidate correlate of consciousness than the P3b, at least under conditions of IB, since it was observed in phase two when the word pairs were clearly perceived.

Comparatively, the P3b was absent in this same phase, which implies it is *not* a neural correlate of consciousness. This absence of P3b adds to a growing body of evidence that suggests the P3b is not a neural correlate of consciousness but is more likely reflective of post-perceptual processes, such as neuro-cognitive functions that act upon visual information after it has already been consciously processed (e.g., Cohen et al., 2020).

It is worth noting that the elicitation of a P3 in the current study is, itself, not trivial. Some previous studies have avoided simultaneous analysis of N400 and P3 responses to the same stimuli due to component overlap (Daltrozzo et al., 2012; Deacon et al., 2004). For example, targets elicit a P3 due to decision making and stimulus classification processes. If these same stimuli are time-locked for N400 analysis, then the possibility arises that P3 and N400 simultaneously occur, which may suppress one or the other component, or otherwise confound the measurement of either component. In the current study, our experimental stimuli allowed us to feasibly examine both through different contrasts. Namely, the task in phase three required participants to respond to stimuli irrespective of whether words were presented. That is, in addition to word pairings themselves, stimulus sequences with no words at all were targets that required a response. In effect, this allowed for two separate contrasts with respect to an analysis of the target word (i.e., the second word): one between related and unrelated word pairs (to test for the N400) and one between words and non-words (to test for the P3b). Yet while the presence of a P3b did not impact our capacity to detect the N400 (even within the same temporal window), there was indication that the N400 may have obscured the P3b, since negative amplitude differences were apparent within the ERP time window preceding the P3b (see Figure 4E) and the component topographic distribution was more lateralized than is typical, presumably due to the strong negativity over central regions resulting from semantic properties of the word pairs. Even so, P3b occurred in alignment with previous studies (Pitts et al., 2014), implying that both components can be observed to target stimuli in stimulus pair paradigms, so long as stimuli allow for sufficient contrasts for detection/separation of each component.

### Limitations and Future Directions

There are several broad recommendations we can provide for future work in terms of further maximising the chance of detecting an N400 during IB (or electrophysiological evidence for semantic processing more generally). First, to allow for a between-subject comparison of noticers and non-noticers in IB studies, it is generally ideal (although not always practical/feasible) for an approximately equal distribution of noticers and non-noticers during the critical experimental period. In the current study, all participants failed to notice the words in phase one. This meant that we were unable to perform a between-subject analysis in this phase to assess for differences associated with spontaneous awareness of the words. The facilitation of both a within- and between-subject comparison is a strength of previous IB works (e.g., Shafto & Pitts, 2015) that we were therefore unable to reproduce.

More notably, several participants were excluded from our analyses due to failing to notice words even in phase two—that is, after being cued to their presence via the post-phase questionnaire. These issues suggest that our experimental set-up was potentially too taxing, such that the task was so demanding for some participants that they were unable to perceive the word stimuli even after being informed of their presence. It may be worthwhile for future work to reduce the difficulty of the change detection task or possibly increase the salience of the words, since either of these factors might in theory bolster rates of noticing in the present experiment.

The use of different word pairings need also be considered. The choice of stimuli is perhaps the most critical component in any N400 work, given that the N400 is elicited based on the violation of a semantic context established by the experimental stimuli used. The choice of words in other N400 literature centres on the degree to which respondents would supply a given word when prompted with a sentence or context, referred to as *cloze* probability (Kutas & Federmeier, 2011). Here, we used reasonably liberal Wu and Palmer value cut offs for our choice of unrelated and related word pairs. Whilst the strong N400 observed in phase three demonstrates that our stimuli could reliably elicit an N400, the use of word pairs with more stringent cut offs may be a simple method of improving the chance to detect N400 activity in phase one of the current task (at least in principle).

## Conclusion

Through probing the degree to which the N400 is elicited under conditions of IB, the current study provides novel evidence toward the debate on whether semantic processing can occur unconsciously. Our findings constrain the possible scenarios in which an N400 might be elicited in the absence of awareness. Task irrelevant stimuli that are not perceived do not elicit N400 activity. Task relevancy may therefore be an underappreciated factor in determining whether semantic processing can occur for stimuli that are not perceived. Task relevancy might, for example, afford a stimulus some pre-conscious attentional allocation, which subsequently facilitates processing to a stage of semantic categorization, even when the stimulus is not perceived. Task irrelevant stimuli, on the other hand, are afforded no such benefit. What remains unknown is whether the N400 is elicited for unconscious task relevant stimuli. Based on our account, the N400 should be elicited if it is afforded some pre-conscious attentional allocation due to its task relevance. Prior work aside (Luck et al., 1996), only through an orthogonal manipulation of both task relevancy and awareness can this question be directly addressed. The outcome could resolve the question of why semantic processing can occur unconsciously in some instances, but not others.

## Footnotes

1 Noting it is difficult to see how this could be done in practice, since the outcome measure in their work was reliant on the congruency between the two.

2 Given their research aim, participants were excluded if they noticed the words. Nevertheless, rates of IB were exceptionally high.

## Notes

https://osf.io/gpxub/

### Competing Interest Statement

The authors have declared no competing interest.

## References

1. Bae, G. Y., & Luck, S. J. (2018). Dissociable decoding of spatial attention and working memory from EEG oscillations and sustained potentials. The Journal of neuroscience, 38(2), 409–422. https://doi-org.virtual.anu.edu.au/10.1523/JNEUROSCI.2860-17.2017

2. Bae, G. Y., & Luck, S. J. (2019). Decoding motion direction using the topography of sustained ERPs and alpha oscillations. NeuroImage, 184(1), 242–255. 10.1016/j.neuroimage.2018.09.029

3. Batterink, L., Karns, C. M., Yamada, Y., & Neville, H. (2009). The role of awareness in semantic and syntactic processing: An ERP attentional blink study. Journal of Cognitive Neuroscience, 22(11), 2514–2529. doi: 10.1162/jocn.2009.21361

4. Brown, C. & Hagoort, P. (1993). The processing nature of the N400: Evidence from masked priming. Journal of Cognitive Neuroscience, 5(1), 34–44. 10.1162/jocn.1993.5.1.34

5. Damian M. F. (2001). Congruity effects evoked by subliminally presented primes: automaticity rather than semantic processing. Journal of experimental psychology. Human perception and performance, 27(1), 154–165. 10.1037//0096-1523.27.1.154

6. Davis, M. H., Coleman, M. R., Absalom, A. R., Rodd, J. M., Johnsrude, I. S., Matta, B. F., Owen, A. M., & Menon, D. K. (2007). Dissociating speech perception and comprehension at reduced levels of awareness. Proceedings of the National Academy of Sciences of the United States of America, 104(41), 16032–16037. 10.1073/pnas.0701309104

7. Deacon, D., Hewitt, S., Yang, C., & Nagata, M. (2000). Event-related potential indices of semantic priming using masked and unmasked words: Evidence that the N400 does not reflect a post-lexical process. Brain research. Cognitive brain research, 9(2), 137–146. https://doi-org.virtual.anu.edu.au/10.1016/s0926-6410(99)00050-6

8. Deacon, D., & Shelley-Tremblay, J. (2000). How automatically is meaning accessed: a review of the effects of attention on semantic processing. Frontiers in bioscience: a journal and virtual library, 5, E82–E94. https://doi-org.virtual.anu.edu.au/10.2741/deacon

9. Dehaene, S., Naccache, L., Le Clec’H, G., Koechlin, E., Mueller, M., Dehaene-Lambertz, G., van de Moortele, P. F., & Le Bihan, D. (1998). Imaging unconscious semantic priming. Nature, 395(6702), 597–600. 10.1038/26967

10. Erlbeck, H., Kubler, A., Kotchoubey, B., & Veser, S. (2014). Task instructions modulate the attentional mode affecting the auditory MMN and the semantic N400. Frontiers in Human Neuroscience, 8, 654. 10.3389/fnhum.2014.00654

11. Förster, J., Koivisto, M., & Revonsuo, A. (2020). ERP and MEG correlates of visual consciousness: The second decade. Consciousness and Cognition, 80. 10.1016/j.concog.2020.102917

12. Grossi G. (2006). Relatedness proportion effects on masked associative priming: an ERP study. Psychophysiology, 43(1), 21–30. https://doi-org.virtual.anu.edu.au/10.1111/j.1469-8986.2006.00383.x

13. Guzzon, D. & Casco, C. (2011). The effect of visual experience on texture segmentation without awareness. Vision Research, 51, 2509–2516. doi: 10.1016/j.visres.2011.10.006

14. Holcomb, P. J., Reder, L., Misra, M., & Grainger, J. (2005). The effects of prime visibility on ERP measures of masked priming. Brain research. Cognitive brain research, 24(1), 155–172. 10.1016/j.cogbrainres.2005.01.003

15. Hutchinson, B. (2019). Toward a theory of consciousness: A review of the neural correlates of inattentional blindness. Neuroscience and Biobehavioral Reviews, 104, 87–99. 10.1016/j.neubiorev.2019.06.003

16. Hutchinson, B. T., Pammer, K., Bandara, K., & Jack, B. N. (2022). A tale of two theories: A meta-analysis of the attention set and load theories of inattentional blindness. Psychological Bulletin, 148(5-6), 370–396. 10.1037/bul0000371.

17. Hutchinson, B. T., Wilkinson, N., Robertson, G., Budd, A., Nicholls, M. E. R., & Griffiths, O. (2023). An investigation of inattentional blindness using gaze and frequency tagging. Journal of experimental psychology. Human perception and performance, 49(10), 1310–1329. 10.1037/xhp0001143

18. Jarosz, A. F. & Wiley, J. (2014). What are the odds? A practical guide to computing and reporting bayes factor. Journal of Problem Solving, 7. 10.7771/1932-6246.1167

19. Kang, M., Blake, R., & Wooman, G. (2011). Semantic analysis does not occur in the absence of awareness induced by interocular suppression. Journal of Neuroscience, 31(38), 13535–13545. 10.1523/JNEUROSCI.1691-11.2011

20. Kemp, A., Eddins, D., Shrivastav, R., & Wray, A. H. (2019). Effects of task difficulty on neural processes underlying semantics: An event-related potentials study. Journal of Speech, Language and Hearing Research, 62(2), 367–386. 10.1044/2018_JSLHR-H-17-0396

21. Kiefer M. (2002). The N400 is modulated by unconsciously perceived masked words: Further evidence for an automatic spreading activation account of N400 priming effects. Brain research. Cognitive brain research, 13(1), 27–39. https://doi-org.virtual.anu.edu.au/10.1016/s0926-6410(01)00085-4

22. Kouider, S., Gardelle, V., Sackur, J., & Dupoux, E. (2011). How rich is consciousness? The partial awareness hypothesis. Trends in Cognitive Sciences, 14(7), 301–307. 10.1016/j.tics.2010.04.006

23. Kutas, M. & Federmeier, K. (2011). Thirty years and counting: Finding meaning in the N400 component of the event-related brain potential (ERP). Annual Review of Psychology, 62, 621–647. 10.1146/annurev.psych.093008.131123

24. Kutas, M. & Hillyard, S. A. (1980). Reading senseless sentences: Brain potentials reflect semantic incongruity. Science, 207, 203–205. doi: 10.1126/science.7350657

25. Luck, S. J., Vogel, E. K., & Shapiro, K. L. (1996). Word meanings can be accessed but not reported during the attentional blink. Nature, 383(6601), 616–618. https://doi-org.virtual.anu.edu.au/10.1038/383616a0

26. Matsuyoshi, D., Ikeda, T., Sawamoto, N., Kakigi, R., Fukuama, H., & Osaka, N. (2010). Task-irrelevant memory load induces inattentional blindness without temporo-parietal suppression. Neuropsychologia, 48(10), 3094–3101. doi: 10.1016/j.neuropsychologia.2010.06.021

27. McCarthy, G., & Nobre, A. C. (1993). Modulation of semantic processing by spatial selective attention. Electroencephalography and clinical neurophysiology, 88(3), 210–219. 10.1016/0168-5597(93)90005-a

28. Mongelli, V., Meijs, E. L., Van Gaal, S., & Hagoort, P. (2019). No language unification without neural feedback: how awareness affects sentence processing. Neuroimage, 202:116063. 10.1016/j.neuroimage.2019.116063

29. Nakamura, K., Makuuchi, M., Oga, T., Mizuochi-Endo, T., Iwabuchi, T., Nakajima, Y., & Dehaene, S. (2018). Neural capacity limits during unconscious semantic processing. The European Journal of Neuroscience, 47(8), 929–937. 10.1111/ejn.13890

30. Neumann, M. & Schweinberger, S. (2008). N250r and N400 ERP correlates of immediate famous face repetition are independent of perceptual load. Brain Research, 1239, 181–190. 10.1016/j.brainres.2008.08.039

31. Pitts, M. A., Martínez, A., & Hillyard, S. A. (2012). Visual processing of contour patterns under conditions of inattentional blindness. Journal of Cognitive Neuroscience, 24(2), 287–303. doi: 10.1162/jocn_a_0011

32. Pitts, M. A., Padwal, J., Fennelly, D., Martinez, A., & Hillyard, S. A. (2014). Gamma band activity and the p3 reflect post-perceptual processes, not visual awareness. NeuroImage, 101, 337–350. doi: 10.1016/j.neuroimage.2014.07.024

33. Pugnaghi, G., Memmert, D., & Kreitz, C. (2020). Loads of unconscious processing: The role of perceptual load in processing unattended stimuli during inattentional blindness. Attention, Perception, and Psychophysics, 82(5), 2641–2651. 10.3758/s13414-020-01982-8

34. Relander, K., Rama, P., & Kujala, T. (2008). Word semantics is processed even without attentional effort. Journal of Cognitive Neuroscience, 21(8), 1511–1522. Doi: 10.1162/jocn.2009.21127

35. Ro, T., Singhal, N. S., Breitmeyer, B. G., & Garcia, J. O. (2009). Unconscious processing of color and form in metacontrast masking. Perception, & Psychophysics 71, 95–103. 10.3758/APP.71.1.95

36. Schelonka, K., Graulty, C., Canseco-Gonzalez, E., & Pitts, M. A. (2017). ERP signatures of conscious and unconscious word and letter perception in an inattentional blindness paradigm. Consciousness and Cognition, 54, 56–71. doi: 10.1016/j.concog.2017.04.009

37. Schnuerch, R., Kreitz, C., Gibbons, H., & Memmert, D. (2016). Not quite so blind: Semantic processing despite inattentional blindness. Journal of experimental psychology: Humanperception and performance, 42(4), 459–463. 10.1037/xhp0000205

38. Scholte, H. S., Witteveen, S. C., Spekreijse, H., & Lamme, V. A. F. (2006). The influence of inattention on the neural correlates of scene segmentation. Brain Research, 1076(1), 106–115. doi: 10.1016/j.brainres.2005.10.051

39. Shafto, J. P. & Pitts, M. A. (2015). Neural signatures of conscious face perception in an inattentional blindness paradigm. The Journal of Neuroscience, 35(31), 10940–10948. doi: 10.1523/jneurosci.0145-15.2015

40. Simons, D. J., & Chabris, C. F. (1999). Gorillas in our midst: Sustained inattentional blindness for dynamic events. Perception, 28, 1059–1074. doi: 10.1068/p281059

41. Sklar, A. Y., Levy, N., Goldstein, A., Mandel, R., Maril, A., & Hassin, R. R. (2012). Reading and doing arithmetic nonconsciously. Proceedings of the National Academy of Sciences of the United States of America, 109(48), 19614–19619. 10.1073/pnas.1211645109

42. Stenberg, G., Lindgren, M., Johansson, M., Olsson, A., & Rosén, I. (2000). Semantic processing without conscious identification: Evidence from event-related potentials. Journal of Experimental Psychology: Learning, Memory, and Cognition, 26(4), 973–1004. 10.1037/0278-7393.26.4.973

43. Todd, J. J., Fougnie, D., & Marois, R. (2005). Visual short-term memory load suppresses temporo-parietal junction activity and induces inattentional blindness. Psychological Science, 16(12), 965–972. doi: 10.1111/j.1467-9280.2005.01645.x

44. Treder, M. S. (2020). MVPA-Light: A classification and regression toolbox for multi-dimensional data. Frontiers in neuroscience, 14, 289. 10.3389/fnins.2020.00289

45. van Gaal, S., Naccache, L., Meuwese, J. D., van Loon, A. M., Leighton, A. H., Cohen, L., & Dehaene, S. (2014). Can the meaning of multiple words be integrated unconsciously? Philosophical transactions of the Royal Society of London. Series B, Biological sciences, 369(1641), 20130212. 10.1098/rstb.2013.0212

46. Vandenbroucke, A. R., Fahrenfort, J. J., Sligte, I. G., & Lamme, V. A. (2014). Seeing without knowing: Neural signatures of perceptual inference in the absence of report. Journal of Cognitive Neuroscience, 26(5), 955–969. doi: 10.1162/jocn_a_00530

47. Warin, M., & Volk, H.M. (2004). Using WordNet and semantic similarity to disambiguate an ontology. Technical Report. Stockholms Universitet Institutionen för lingvis

