## supplementary materials for "No Electrophysiological Evidence for Semantic Processing During Inattentional Blindness"

**Word Pairs**

| **Related** | **Word A** | **Word B** | **Relatedness (WUP)** | **Unrelated** | **Word A** | **Word B** | **Relatedness (WUP)** |
| --- | --- | --- | --- | --- | --- | --- | --- |
| 1 | CREAM | SUGAR | 0.7 | 1 | SIREN | SUGAR | 0.32 |
| 2 | DOCTOR | NURSE | 0.88 | 2 | CREEK | NURSE | 0.4 |
| 3 | MOUSE | CHEESE | 0.64 | 3 | BULL | CHEESE | 0.23 |
| 4 | SCHOOL | DESK | 0.6 | 4 | DREAM | DESK | 0.22 |
| 5 | TURTLE | SHELL | 0.67 | 5 | MAIL | SHELL | 0.4 |
| 6 | ROOF | HOUSE | 0.67 | 6 | CARAMEL | HOUSE | 0.3 |
| 7 | PAGE | BOOK | 0.63 | 7 | JELLY | BOOK | 0.4 |
| 8 | FLAKE | SNOW | 0.67 | 8 | SURGEON | SNOW | 0.27 |
| 9 | ANKLE | FOOT | 0.67 | 9 | SEED | FOOT | 0.29 |
| 10 | CIRCLE | SQUARE | 0.83 | 10 | POPPY | SQUARE | 0.17 |
| 11 | EYES | NOSE | 0.84 | 11 | COOKIE | NOSE | 0.32 |
| 12 | HORSE | BARN | 0.7 | 12 | LIQUID | BARN | 0.38 |
| 13 | SHOWER | SOAP | 0.8 | 13 | SNAIL | SOAP | 0.26 |
| 14 | GLASS | BOTTLE | 0.86 | 14 | TENNIS | BOTTLE | 0.19 |
| 15 | BABY | ADULT | 0.83 | 15 | PENNY | ADULT | 0.27 |
| 16 | CALF | COWS | 0.67 | 16 | COAST | COWS | 0.31 |
| 17 | MILK | CEREAL | 0.88 | 17 | CITY | CEREAL | 0.36 |
| 18 | FORK | KNIFE | 0.78 | 18 | POSTER | KNIFE | 0.24 |
| 19 | FLOOR | CARPET | 0.71 | 19 | TIDE | CARPET | 0.25 |
| 20 | LEAF | TWIG | 0.8 | 20 | SPACE | TWIG | 0.25 |
| 21 | RIVER | WATER | 0.83 | 21 | TRUNK | WATER | 0.33 |
| 22 | PRISON | JAIL | 0.92 | 22 | SQUID | JAIL | 0.4 |
| 23 | DRINK | JUICE | 0.8 | 23 | NIGHT | JUICE | 0.29 |
| 24 | BURN | FIRE | 1 | 24 | BONE | FIRE | 0.38 |
| 25 | CAKE | PIES | 0.82 | 25 | DESERT | PIES | 0.33 |
| 26 | TREE | BRANCH | 0.75 | 26 | QUEEN | BRANCH | 0.21 |
| 27 | LIPS | TEETH | 0.75 | 27 | TIGHTS | TEETH | 0.22 |
| 28 | BIKE | WHEEL | 1 | 28 | CORN | WHEEL | 0.38 |
| 29 | APPLE | ORANGE | 0.88 | 29 | JOKE | ORANGE | 0.38 |
| 30 | SWEETS | CANDY | 0.95 | 30 | CLONE | CANDY | 0.35 |
| 31 | HAIR | COMB | 0.82 | 31 | HEART | COMB | 0.21 |
| 32 | BREAD | TOAST | 0.94 | 32 | COMPASS | TOAST | 0.3 |
| 33 | HAMMER | NAIL | 0.73 | 33 | BREEZE | NAIL | 0.3 |
| 34 | BOYS | GIRLS | 0.85 | 34 | WEEK | GIRLS | 0.29 |
| 35 | LOVE | HATE | 0.88 | 35 | ROOM | HATE | 0.24 |
| 36 | LEMON | LIME | 0.93 | 36 | PARTY | LIME | 0.22 |
| 37 | STAR | PLANET | 0.88 | 37 | GAME | PLANET | 0.25 |
| 38 | CROWN | KING | 0.67 | 38 | MONTH | KING | 0.29 |
| 39 | LAKE | POOL | 0.92 | 39 | WORD | POOL | 0.35 |
| 40 | PAINT | BRUSH | 0.7 | 40 | TEAM | BRUSH | 0.24 |
| 41 | LEGS | KNEE | 0.89 | 41 | PARENT | KNEE | 0.33 |
| 42 | BRAIN | SKULL | 0.6 | 42 | SUIT | SKULL | 0.29 |
| 43 | SONG | MUSIC | 0.89 | 43 | WOOD | MUSIC | 0.4 |
| 44 | WALK | HIKE | 0.96 | 44 | BELL | HIKE | 0.18 |
| 45 | FINGER | HAND | 0.89 | 45 | STREET | HAND | 0.32 |
| 46 | PHONE | CALL | 1 | 46 | SAND | CALL | 0.3 |
| 47 | DINNER | LUNCH | 0.89 | 47 | ISLAND | LUNCH | 0.4 |
| 48 | RICE | WHEAT | 0.93 | 48 | SEAL | WHEAT | 0.4 |
| 49 | SOCK | SHOE | 0.7 | 49 | MEAT | SHOE | 0.38 |
| 50 | LOCK | KEYS | 0.86 | 50 | SOCCER | KEYS | 0.19 |
| 51 | BIRD | WING | 0.67 | 51 | BEARD | WING | 0.32 |
| 52 | PISTOL | BULLET | 0.8 | 52 | TAIL | BULLET | 0.3 |
| 53 | CLOUD | RAIN | 0.8 | 53 | YARN | RAIN | 0.33 |
| 54 | SUMMER | WINTER | 0.88 | 54 | CAMEL | WINTER | 0.17 |
| 55 | PLANE | TRAVEL | 0.67 | 55 | DAISY | TRAVEL | 0.19 |
| 56 | MUSEUM | GALLERY | 0.67 | 56 | SOIL | GALLERY | 0.35 |
| 57 | BEER | WINE | 0.84 | 57 | ENGINE | WINE | 0.3 |
| 58 | SINK | TOWEL | 0.67 | 58 | PEEL | TOWEL | 0.38 |
| 59 | GIFT | PRESENT | 1 | 59 | BOWL | PRESENT | 0.21 |
| 60 | PILLOW | BLANKET | 0.71 | 60 | SHADOW | BLANKET | 0.2 |
| 61 | WINDOW | DRAPES | 0.75 | 61 | PIZZA | DRAPES | 0.33 |
| 62 | PENCIL | ERASER | 0.84 | 62 | FROWN | ERASER | 0.24 |
| 63 | METER | INCH | 0.82 | 63 | BANK | INCH | 0.35 |
| 64 | GLOVES | MITTEN | 0.96 | 64 | VOICE | MITTEN | 0.21 |
| 65 | COUGH | SNEEZE | 0.89 | 65 | LETTUCE | SNEEZE | 0.33 |
| 66 | GHOST | MONSTER | 0.7 | 66 | BLUSH | MONSTER | 0.38 |
| 67 | MOON | PHASE | 0.62 | 67 | SODA | PHASE | 0.35 |
| 68 | WITCH | WIZARD | 0.86 | 68 | KISS | WIZARD | 0.27 |
| 69 | MOUTH | TONGUE | 0.71 | 69 | YEAR | TONGUE | 0.27 |
| 70 | WASP | HORNET | 0.93 | 70 | CRIB | HORNET | 0.4 |
| 71 | HONEY | BEES | 0.61 | 71 | BUTTER | BEES | 0.3 |
| 72 | KETCHUP | MUSTARD | 0.91 | 72 | ANGER | MUSTARD | 0.22 |
| 73 | FLOWER | PETAL | 0.76 | 73 | HORROR | PETAL | 0.27 |
| 74 | FATHER | MOTHER | 1 | 74 | COMEDY | MOTHER | 0.21 |
| 75 | SISTER | BROTHER | 0.92 | 75 | TREAT | BROTHER | 0.32 |
| 76 | PEACH | PLUM | 0.93 | 76 | BLOOD | PLUM | 0.27 |
| 77 | CROW | RAVEN | 0.93 | 77 | BALLET | RAVEN | 0.18 |
| 78 | FISH | SHARK | 0.91 | 78 | LUCK | SHARK | 0.29 |
| 79 | CHICKEN | ROOSTER | 0.97 | 79 | TRUTH | ROOSTER | 0.17 |
| 80 | MOVIE | FILM | 1 | 80 | FEAR | FILM | 0.38 |
| 81 | PORK | BEEF | 0.88 | 81 | DANCE | BEEF | 0.36 |
| 82 | GUITAR | DRUMS | 0.82 | 82 | SLEEP | DRUMS | 0.21 |
| 83 | PLANT | CACTUS | 0.84 | 83 | PEACE | CACTUS | 0.22 |
| 84 | LETTER | STAMP | 0.8 | 84 | CELERY | STAMP | 0.22 |
| 85 | SWEATER | COAT | 0.86 | 85 | THRILL | COAT | 0.2 |
| 86 | BULB | LAMP | 0.91 | 86 | SHEEP | LAMP | 0.37 |
| 87 | MONEY | COIN | 0.84 | 87 | BANANA | COIN | 0.21 |
| 88 | MEAL | FOOD | 0.86 | 88 | CELLO | FOOD | 0.33 |
| 89 | TRACK | TRAIN | 0.75 | 89 | BACON | TRAIN | 0.3 |
| 90 | SHIRT | PANTS | 0.9 | 90 | PAPER | PANTS | 0.3 |
| 91 | SCARF | NECK | 0.67 | 91 | BRICK | NECK | 0.38 |
| 92 | NEEDLE | THREAD | 0.76 | 92 | CABBAGE | THREAD | 0.32 |
| 93 | SOFA | COUCH | 1 | 93 | BADGE | COUCH | 0.26 |
| 94 | WATCH | CLOCK | 0.92 | 94 | VINEGAR | CLOCK | 0.26 |
| 95 | CAMP | TENT | 1 | 95 | SMILE | TENT | 0.29 |
| 96 | JEANS | DENIM | 1 | 96 | SALT | DENIM | 0.35 |
| 97 | FIDDLE | VIOLIN | 1 | 97 | ADVICE | VIOLIN | 0.21 |
| 98 | TRUMPET | TUBA | 0.92 | 98 | HAIL | TUBA | 0.27 |
| 99 | BOAT | SHIP | 0.92 | 99 | LUNG | SHIP | 0.27 |
| 100 | TUNA | SALMON | 0.88 | 100 | TRAFFIC | SALMON | 0.4 |
